# Biochemical and mathematical lessons from the evolution of the SARS-CoV-2 virus: paths for novel antiviral warfare

**DOI:** 10.1101/2020.07.31.230607

**Authors:** Nicolas Cluzel, Amaury Lambert, Yvon Maday, Gabriel Turinici, Antoine Danchin

## Abstract

In the fight against the spread of COVID-19 the emphasis is on vaccination or on reactivating existing drugs used for other purposes. The tight links that necessarily exist between the virus as it multiplies and the metabolism of its host are systematically ignored. Here we show that the metabolism of all cells is coordinated by the availability of a core building block of the cell’s genome, cytidine triphosphate (CTP). This metabolite is also the key to the synthesis of the viral envelope and to the translation of its genome into proteins. This unique role explains why evolution has led to the early emergence in animals of an antiviral immunity enzyme, viperin, that synthesizes a toxic analogue of CTP. The constraints arising from this dependency guide the evolution of the virus. With this in mind, we explored the real-time experiment taking place before our eyes using probabilistic modelling approaches to the molecular evolution of the virus. We have thus followed, almost on a daily basis, the evolution of the composition of the viral genome to link it to the progeny produced over time, particularly in the form of blooms that sparked a firework of viral mutations. Some of those certainly increase the propagation of the virus. This led us to make out the critical role in this evolution of several proteins of the virus, such as its nucleocapsid N, and more generally to begin to understand how the virus ties up the host metabolism to its own benefit. A way for the virus to escape CTP-dependent control in cells would be to infect cells that are not expected to grow, such as neurons. This may account for unexpected body sites of viral development in the present epidemic.

## Introduction

The development of the COVID-19 pandemic is being explored in a myriad of articles. Despite this abundance, and because of our anthropocentrism, it is exceptional that these studies focus on the virus’ standpoint. Of course, much work is looking into the details of the composition and structure of the SARS-CoV-2 virus genome, the proteins it codes for and its animal-infecting relatives. However, there are very few major studies on how the virus exploits the metabolism of its host’s cells. The urgent necessity to contain the disease led investigators to emphasize vaccination or, more generally, the involvement of the host’s immune system. It is well known, alas, that while it has sometimes been relatively easy to generate a vaccine that is both effective and harmless against a widespread disease, the opposite is also true. There are still very serious and very common diseases for which there is no vaccination. Vaccinating effectively assumes, in particular, that the progeny of a pathogen remains the same long enough to prevent escape of the immune response triggered by the vaccine. Coronaviruses are viruses made up of a long genome and an envelope. The length of the genome could have led to a very high mutation rate, but these viruses, thus avoiding the universal constraint of Muller’s ratchet - see **Box** - have recruited a specific function that proofreads and corrects replication errors [1]. This means that, while coronaviruses do indeed tend to produce genetic variants over time, the number of these variants remains quite low. This mutation rate may appear very limited, but the sheer number of viral particles generated during an infection is enormous, while the human population currently recognized as infected exceeds twenty million people. It follows that the mutation rate per nucleotide - of course very heterogeneous due to the selection pressure on certain locations in the genome - is around 8 x 10^-4^ changes per site per year [2].

Here, this situation was placed in the perspective of the fundamental theorem of natural selection proposed by Fisher, which links the evolution of environmental fitness and genetic variance [3]. We wished to use the marks left by the evolution of the virus’ fitness - observed in the form of genomic sequences - in the presence of the biochemical constraints that bias the choices available for evolution. We had to take into account, however, that the terms of the problem are not as explicit as one might have wished: fitness is not known, nor are the time markers (estimated from phylogenetic trees or simply taken as physical time) and the frequency of certain strains in the phylogenetic trees may be less due to natural selection than to heterogeneity in sampling and sequencing depth. This motivated our use of procedures that are robust enough to cope with these uncertainties. Nevertheless, the advantage of such an analysis is that it allowed us to propose anticipations for the evolution of the virus. It is therefore an explicit means of feeding epidemiological or clinical models with relevant observations.

In this context, it seemed to us of great interest to explore the details of how SARS-CoV-2 mutated over time, in the various places where COVID-19 has spread, highlighting relevant descents in relation with the host metabolism. This should allow us to anticipate some of the future of the virus’ progeny, with important consequences for control of the disease. The analysis of the constraints that govern access to the metabolism of the nucleotides that make up the virus genome has shown us that the content of cytosine (C) in its genome is subjected to strong negative pressure, leading to systematic depletion, over time, in cytosine monophosphate [4]. This bias has long been believed to result from a major causal effect of the “editing” of the C content of the genome by the family of APOBEC deaminating enzymes [5,6]. We now know that it is the organization of the metabolism of pyrimidines in animal cells, and more particularly of cytosine triphosphate [7], which drives the corresponding pressure on evolution (**Figure 1**).

**Figure 1.**
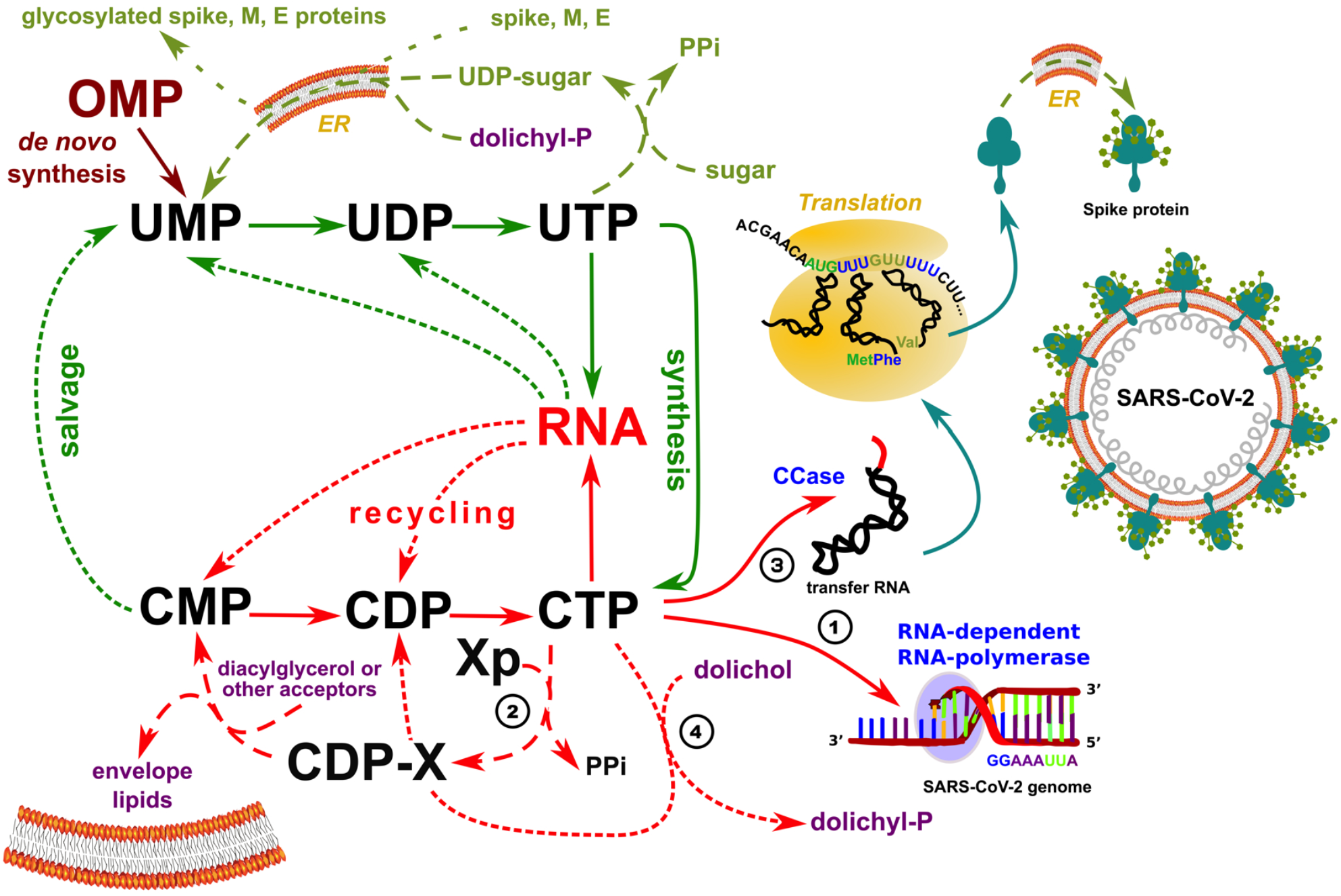
CTP controls all crucial metabolic steps required to build up a functional SARS-CoV-2 virus. 1/ CTP is a precursor of the virus genome; 2/ the lipids of its envelope derive from cytosine-based liponucleotide precursors; 3/ all transfer RNA molecules produced by the host must be maturated to a form ending in a CCA triplet at their 3’OH end; and 4/ post-translational glycosylation of viral proteins, in particular its spike protein require a dolichyl-phosphate anchor in the endoplasmic reticulum (ER) and dolichol kinase is specifically dependent on CTP. See text and reference [7] for details.

Indeed, due to the extreme asymmetry of the replication of the virus - which replicates 50 to 100 times from its complementary template [8] - a genome editing effect of these highly contextdependent enzymes would only be significant when a C into U is modified on the negative RNA template, which would lead to a major enrichment in A of the viral genome, or possibly from a U → C transition due to another class of deaminating enzymes acting on double stranded RNA, ADAR, that deaminates adenine into inosine [9]. Furthermore, both APOBEC and ADAR are highly specific enzymes and this hardly fits with the widespread C → U transitions that we keep observing as the virus evolves. Here, we have focused on the dynamics of the loss of C in the genome, and sought for the locations and the causes of changes in this driving force. In the first paragraph, we summarized the metabolic reasons accounting for this remarkable phenomenon. Subsequently, in the body of the article, we showed that the constraint on the C content of the genome leads to specific descents which can be used to reveal the existence of important functions of the virus as well as the role of the host’s response.

### A universal metabolic requisite, the biosynthesis of cytidine triphosphate (CTP), guides the evolution of the virus

What do we know about the synthesis of the building blocks that allow the generation of a viral particle (a virion)? During a viral infection cells usually stop multiplying. All their resources are quickly diverted in favour of the multiplication of the virus. Yet, growth is a universal property of life. This means that, almost always - differentiated neurons are an exception - the cell’s metabolism that the virus faces is organized to allow cell growth as soon as the opportunity to multiply arises. The moment it infects a cell - again, with the exception of those that do not multiply - any virus will therefore have to manage the metabolic pressure that organizes the availability of the building blocks necessary for its construction. In our usual physical space (three-dimensional), growing introduces an inevitable constraint. The cell must put together the growth of its cytoplasm (threedimensional, therefore), that of the membrane that encloses it (two-dimensional) and that of its genome (one-dimensional, because nucleic acids are linear polymers). However, it is a common metabolism, developed mainly in the cytoplasm, which produces the building materials needed to build up these three major compartments. So, here we have a question similar to the one asked by economists when they raise the question of “non-homothetic” growth [10]. Unfortunately, because life developed from a primitive metabolism in several stages over 3.5 billion years [10], we might fear that many organisms had found an idiosyncratic solution to this constraint, as often witnessed in the huge diversity of life forms. Unexpectedly, it appears that the solution to this quandary is universal: a single metabolite, the nucleotide cytidine triphosphate (CTP), has been recruited to this purpose [4,7].

The key role of CTP appears in four essential places in cellular metabolism, and these places are essential for the formation of new virions. 1/ It is the immediate precursor of one of the four nucleotides forming the genome of the virus; 2/ CTP is required for the synthesis of liponucleotide precursors of the viral envelope; 3/ human transfer RNAs are synthesized from 415 genes which do not encode their 3’OH-CCA terminal end - this sequence is synthesized from CTP by a specific nucleotidyltransferase [12]; and finally 4/ the “decoration” of proteins by complex glycosylations is performed in parallel with their translation in the endoplasmic reticulum (ER) *via* the anchoring of substrates by dolichyl-phosphate, produced by a kinase which uses CTP, not ATP, as its phosphate donor [13]. In addition, intermediate metabolism is based on an original organization of the metabolism of pyrimidines, which systematically recycles and salvages them *via* uridine triphosphate (UTP) which makes CTP a pivot metabolite and limits considerably its availability (**Figure 1**). As a result, accidental replication errors will tend to replace cytosine with uracil in the genome.

### General evolution of the SARS-CoV-2 virus

Using the available sequence data gathered in the SARS-CoV-2 GISAID database (https://www.gisaid.org) we have, like others [14,15], reconstituted a phylogenetic tree of the evolution of the virus. As the sequences of each viral genome, as well as the date of identification of these sequences are known with fairly great precision, this tree makes it possible to explore the orderly lineage of the mutations which appear over time. In particular, unless we can suspect a recombination event due to the infection of the same patient by two or more viruses, when two identical mutations appear in separate branches of the tree, we can assume that this is the result of evolutionary convergence [16]. The reasons for this convergence are discussed on a case by case basis when analysing each relevant mutation. A second observation, which needs to be put in perspective (see below), is that the shape of the tree is not at all homogeneous. We noticed indeed the presence of “blooms” where, at a particular node of the tree, a large number of branches appear, demonstrating an “explosive” appearance of new mutations (**Figure 2**). We have therefore devised a statistical approach that allowed us to characterize them explicitly.

**Figure 2.**
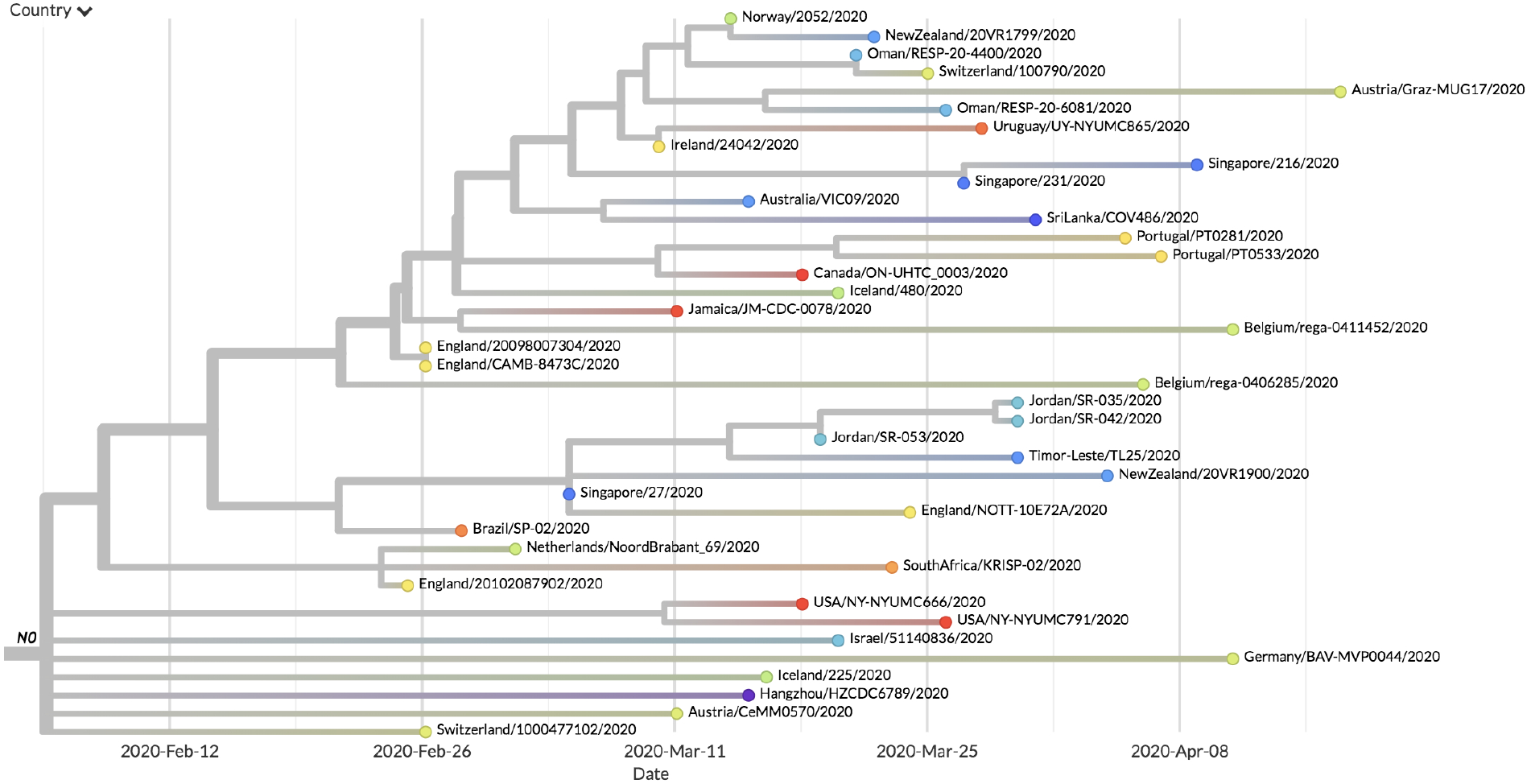
An example of bloom detected by our statistical approach. At node N0, there are 25 different states in the 40 samples of the subtree and a high number of branches. This behaviour differs significantly from that of the other sub-trees.

The causes of these blooms are multiple, but the adaptation of important viral functions can be at their origin, and we retained a few cases of this kind for further discussion (see **Materials and Methods** for the statistical definition of blooms).

### Description and analysis of the evolution of the C content of the genome

Generally speaking, the coronavirus genome tends to evolve by adapting its C content to the metabolism of its host. More specifically SARS-CoV-2 evolves towards forms less rich in C as the epidemic develops [7]. However, this development is not homogeneous.

In the two data sets of interest, 77% of the transitions between pyrimidines are represented by transitions from cytosine to uracil. These transitions represent 48% of all substitutions identified in the first set (respectively, 49% in the second). An important imbalance can also be noted at the level of the transversions, knowing that more than 73% of those pertain to a substitution from purine to pyrimidine in the first set (respectively, 74%). However, only 20% of these 73% lead to the occurrnce of cytosine (respectively, 17%), indicating once again a tendency to favour the generation of uracil, thus demonstrating that the major constraint of the mutagenic process is the availability of each one of the nucleoside triphosphates in the cell. This inhomogeneity is also salient at the tree level. At the level of branch B4 (20% of the samples), the tendency is strongly marked to lose less C as compared to the rest of the tree (**Figure 3**).

**Figure 3:**
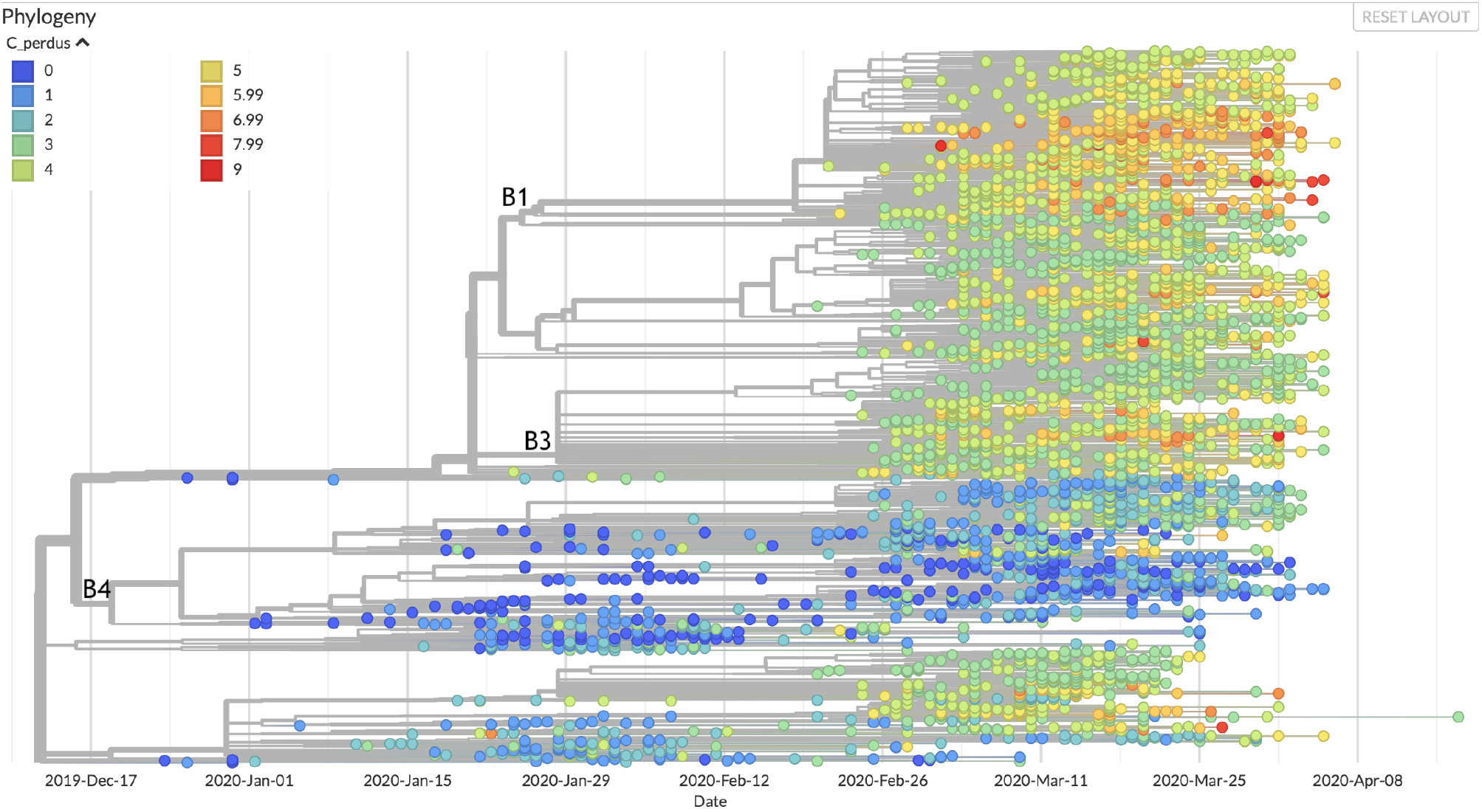
Heat map of C losses from the original sequence. Branches 1 and 4 can be readily discriminated by their extreme values.

Interestingly, this branch is also the one that comprises on average the strains with the least divergence from the original strain of the virus. By contrast, in branch B1, the loss of C looks larger. The rate of virus mutation also seems to be accelerating in this branch, with a rate of transversions 20% higher than the rest of the tree (and also higher transition rates, but in more anecdotal proportions). Finally, for branch B3, the main site of blooms, a 29% decrease in the transition rate of pyrimidines and a 30% decrease in the rate of purines compared to the rest of the tree is noteworthy.

This inhomogeneity can be the consequence of many constraints:

1. The very structure of the genome, which must fold into a compact capsid envelope requires certain regions to maintain the presence of specific C residues. This is the case of the regions which control the origin of replication [8] or transcription, AACGAAC, for example [17]. In the case of the translated regions, the pressure on the presence of C varies depending on its position in the codon trinucleotides. When C is located at the first position of a codon, it is used to input arginine, glutamine, histidine, leucine or proline into proteins. Histidine and glutamine are coded in two codon families, discussed below. For arginine, the selection pressure is lower because the CGN codons can be replaced by AGR codons - we used here the IUPAC convention for labelling nucleotides or aminoacids, e.g. N is for aNy, R for puRine, etc. (https://www.bioinformatics.org/sms/iupac.html). The selection pressure on the leucine content is also lower, since in addition to the CUN codons, this amino acid can be input using the UUR codons. In the second codon position, C is again used to code for proline, but also threonine (ACN), alanine (GCN) and serine (UCN). Again, the latter amino acid escapes a large part of the constraint imposed by the availability of C because it can also use the AGY codons. Finally, the third position of the codons is much less constrained because it can be replaced by U but also by A or G in the families with four codons (alanine, proline, threonine, valine). The two codon families UGY, AGY and NAY are discriminated along a pyrimidine / purine axis. A pyrimidine is used to maintain the same nature of the coded residue, as the codon uses a U or C as the 3 ‘end (aspartate, asparagine, cysteine, histidine and tyrosine). Finally, isoleucine is coded by three codons (AUH), and ending in U or C is taken into account by relevant tRNAs [18];
2. The function of the virus proteins can impose the presence of certain amino acids in their sequence. For example, the proline residue encoded by the CCN codons is not strictly an amino acid, but is essential for the folding of key domains of viral proteins [19];
3. Further stressing the importance of CTP, during evolution, innate antiviral immunity recruited the activity of an enzyme, viperin, which modifies CTP into a form toxic to the development of the virus, 3’-deoxy-3’,4’-didehydro-CTP (ddhCTP) [20]. An interesting consequence of this pathway is that decreasing the C content of the genome will allow the virus replication process to be less sensitive to the presence of this nucleobase. It follows that, during the transfer of a virus relatively rich in C from an animal host to human beings, the evolution towards the loss of C may be transiently concomitant with an increase in its pathogenicity. In the long term, however, the loss of C severely restricts the evolutionary landscape of the virus and most likely will tend to its attenuation [21].

### Examples of correlations allowing us to propose a function for viral proteins

Thousands of mutations have been identified at this date. It is possible to follow their emergence along the tree of its phylogenetic evolution of the virus and then highlight some interesting features that may allow us to anticipate some of its future.

#### Mutations leading to an early translation termination

Mutations leading to premature termination of the virus protein synthesis are expected to appear with high frequency. In the present context, this is all the more likely because the translation termination codons UAA, UAG and UGA do not contain C, and are therefore favoured by the disappearance of this nucleotide. Since most of these mutations lead to non-functional polypeptides, it is generally probable that the affected viruses do not give rise to a significant progeny. It follows that when these mutations are observed - and that they do not result from sequencing errors - they indicate that the role of the truncated protein corresponds to a function which is not critical, or that the protein has remained functional at a sufficient level to allow virus reproduction. However, a few observations allowed us to offer an explanation for the fact that the viruses in question may have survived. Here are three examples which reveal interesting features of the virus.

##### Example 1

In a strain from Iceland, the succession of mutations G1440A (Gly392Asp, protein Nsp2) and G2891A (Ala876Thr, ubiquitin-like domain of protein Nsp3) is now present in multiple world locations [22]. This sequence ends up with C27661U (which modifies amino acid Gln90 into a premature translation end, near the carboxy-terminal end of protein Orf7a). This viral protein is found in the endoplasmic reticulum, the Golgi apparatus and the perinuclear space [23]. Several variants have been identified in the course of the epidemic [24]. Remarkably, several deletions have been isolated in the gene, which suggests that the function of this region is not essential [25]. However, we noticed that many of these mutations, as the one discussed here, keep the small hydrophobic protein Orf7b gene intact, downstream of Orf7a. This very small protein is present in the Golgi apparatus and is also found in the purified virus [26]. It must be noticed that it is synthesized in vivo *via* a frameshift that spans the termination codon of the Orf7a frame (…GAA TGA TT… becomes …GA ATG ATT…). This can be interpreted as a conflict in this region between translation of Orf7a and Orf7b, creating a cost / benefit dilemma for the expression of either one of these proteins. Hence it will be important to monitor the future descent of the virus in this region as it may result in interesting attenuated forms.

##### Example 2

Another succession of mutations that leads to premature translation termination of a viral protein begins with G11083U (protein Nsp6, Leu37Phe). This mutation is now widely distributed worldwide. It is likely to induce a more stable binding of the protein to the ER, possibly favouring coronavirus infection by compromising delivery of viral components to lysosomes for degradation [27]; then we have G1397A (Nsp2, Val378Ile), also likely to favour virus propagation [28]; followed by G29742U (3’UTR of the virus), and U28688C (synonymous); subsequently, we have the couple of mutations C884U (Nsp2 again, Arg207Cys [28]) and G8653U (Nsp4, essential for envelope assembly [29]. The corresponding change (Met2796Ile) is located at the border of the ER lumenal domain of the protein. It is known that, in order to function properly, the ER requires the presence of oxygen [30], and reactive oxygen species (ROS) are associated to misfolding of proteins in this compartment. Nsp4 has a number of cysteine residues, prone to be oxidized. The role of methionine in the parent might be to act as a buffer against ROS, so that the mutant would be slightly attenuated). These mutations are followed by A19073G (in the methylase domain of protein Nsp14, Asp1869Gly, a position that already evolved from SARS-CoV-1 [31], hence likely to be more or less neutral), then the couple with the mutation resulting in end of translation: G27915U, Gly8 to end of translation at the N-terminus of Orf8 and C29077U (synonymous); the succession ends with the couple of mutations leading to synonymous changes C19186U and G23608U. This region of SARS-related coronaviruses is hypervariable. It changes during the course of epidemics, showing that it is subject to ongoing selection pressure, sometimes producing two peptides Orf8a and Orf8b [32]. It corresponds to proteins expressed at the end of the infection cycle. It will be important to monitor the way they function in the course of the evolution of virulence of the virus. This displays a branching that appeared in four different countries and in seven samples, spanning six weeks between the first and the last mutation.

##### Example 3

Here we have a succession of mutations that begin within the 5’end of the virus genome, C241U, followed by mutation C14408U (Pro314Leu) at the end of a zinc finger in replicase Nsp12, which appears in many branches of the evolution tree of the virus. It is discussed in details below (origin of blooms). This mutation is followed by A23403G (Asp614Gly) a widely spread mutation of the spike protein (also discussed below), C3037U (synonymous), mutation G25563U (Gln57His) in Orf3a forming potassium channels is supposed to negatively interfere with the function of the protein [33], C1059U (Thr265Ile) in protein Nsp2, discussed previously, and the triplet G4181A (Ala1305Thr) in the SUD-N domain of protease Nsp3, then mutations G4285U (Glu1340Asp), and G28209U which results in an end of translation at glutamate 106 of protein Orf8. As discussed previously, many mutations, including deletions in Orf8, were frequently observed. This is again an indication that evolution of this regions should be carefully monitored to look for attenuated forms of the virus. This particular mutation to an end of translation is significant as it was found in a sample from Croatia, another one from Thailand, on two significantly separated branches and with one month difference. The sequence of mutations here corresponds to the Thailand sample.

#### Reversal of the tendency of the viral genome to lose its cytosine residues

We have here retained two examples of a situation where, from an upstream branching point in the evolution tree, it appears that the descendants of the virus stop losing their cytosines, and may even tend to regain them. These examples are as follows (**Figure 4**).

**Figure 4.**
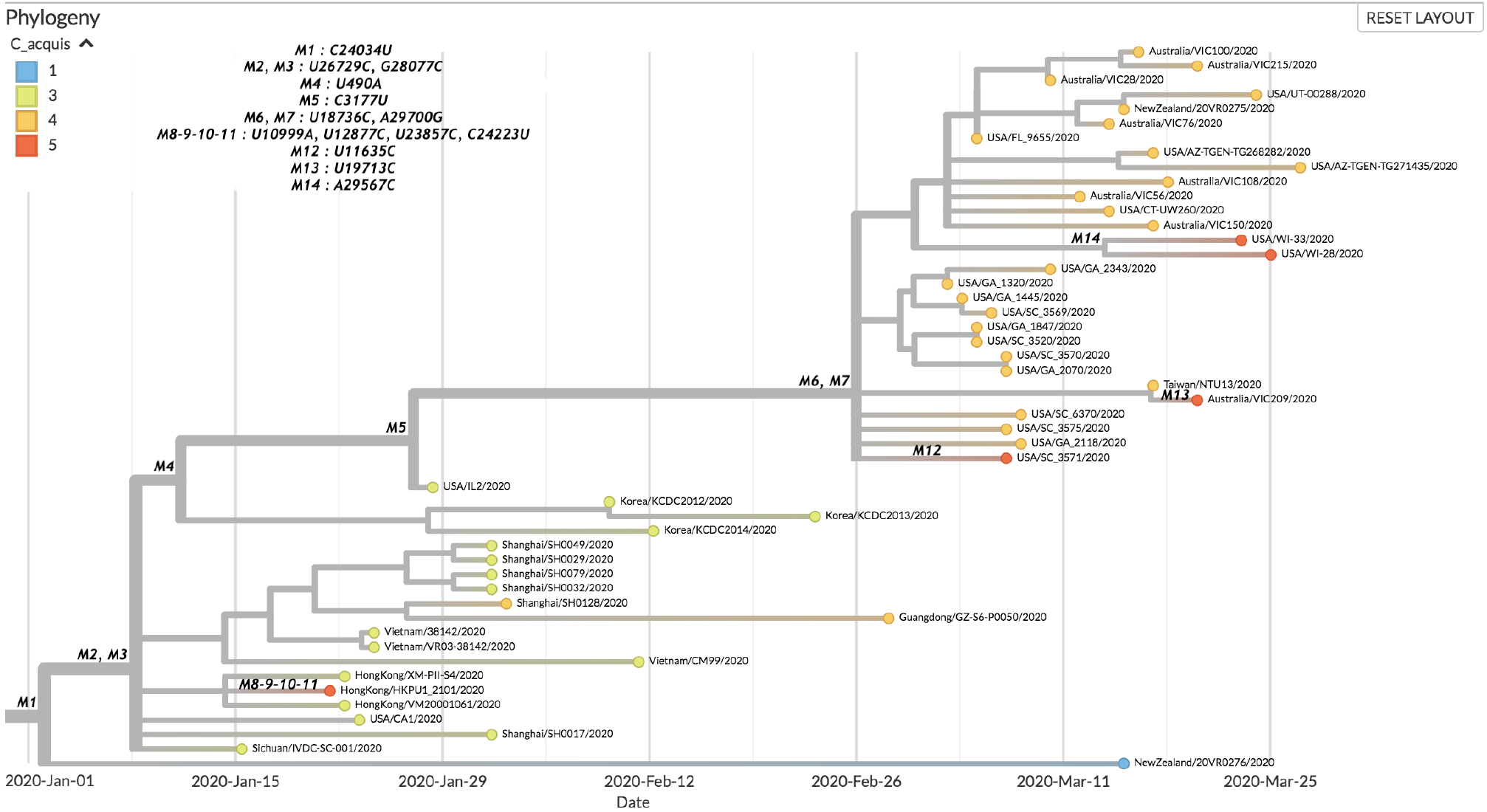
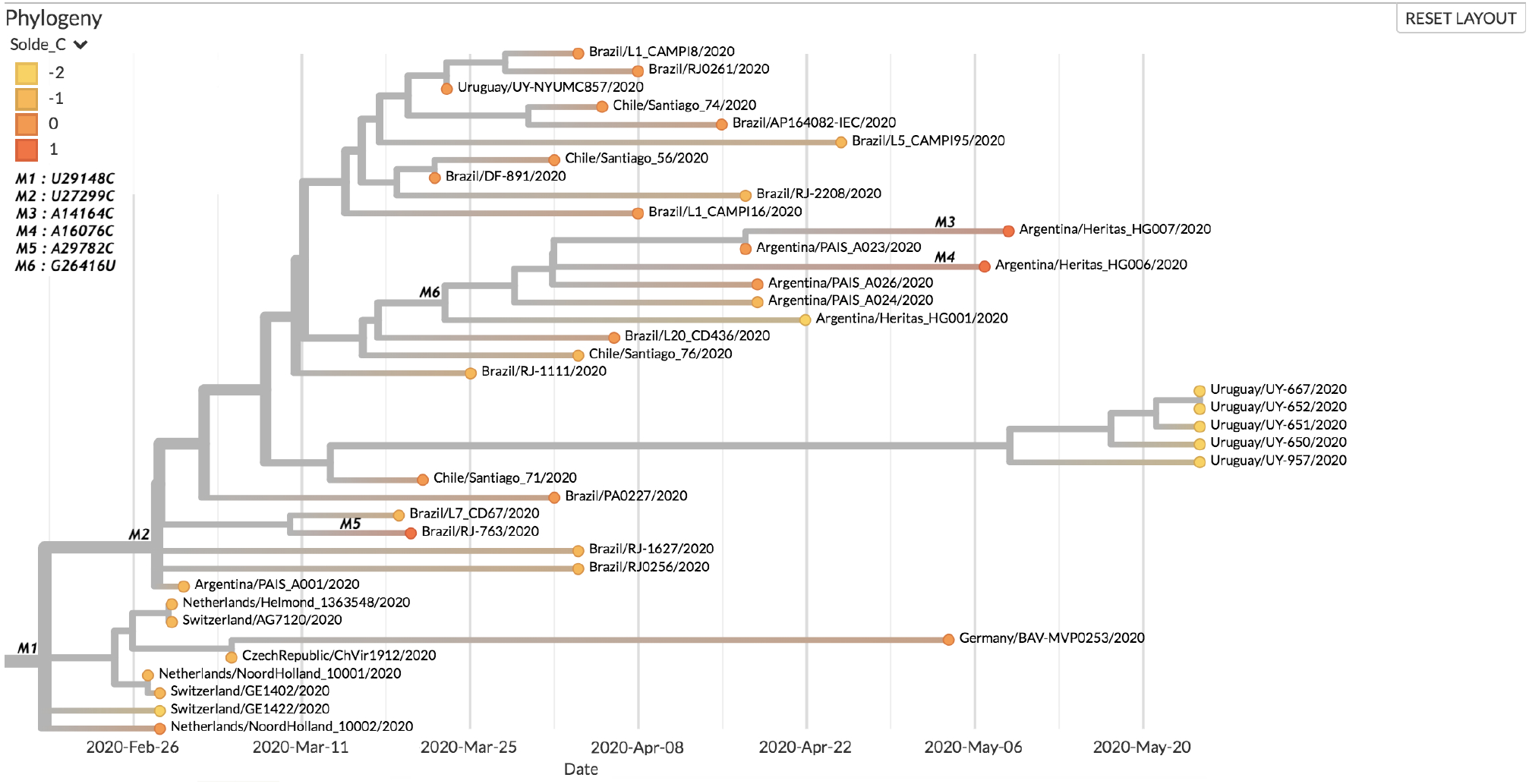
Two sub-trees in the first dataset where the tendency of the genome to lose its cytosine residues is reversed. **Upper panel. First sub-tree.** The sub-samples displayed are those that have acquired the most C, apart from a few isolated samples on other branches. The node with the M1 mutation directly follows those respectively associated with the C8782U and U28144C mutations. **Lower panel. Second sub-tree.** This tree contains a majority of strains with a neutral C balance (both gained and lost), as well as 3 strains with more C gained than lost.

In dataset 1, there are two sub-trees, the first of which is more of an Asian sub-tree with the root of the node associated with the M2 and M3 mutations. The second contains samples from North America and Oceania, and its root node is related to the M6 and M7 mutations. The first tree arises from the succession of C8782U (synonymous), U28144C (Leu84Ser) mutations in the Orf8 protein, whose function was discussed above. It defines a major clade of variants of the virus [24], C24034U (synonymous), and finally the doublet U26729C (synonymous), G28077C (Val62Leu), in the Orf8 protein again. As this is the origin of the observed phenomenon, we are led to believe that it is the alteration of the role of Orf8 (8a or 8b) that is responsible. The Orf8 region is particularly variable and has been clearly implicated in interspecies transmission [34]. A common hypothesis is that the alteration of this gene corresponds to a loss of active function in chiropteran ancestors [35]. Since these are generally richer in cytosine than the human forms [21], one might ask whether one of the functions of this protein is to modulate the activity of CTP synthase.

In fact, the second branch comes from the same descent, to which is added the U490A (Asp75Glu) mutation in the Nsp1 protein, which controls the specific translation of viral RNA [36], systematically associated with the mutation C3177U (Pro971Leu) in the acidic domain, without any clearly identified function, of the multifunctional protease Nsp3 [37], and finally the U18736C (Phe1757Leu) doublet of the exonuclease, N7-methyltransferase Nsp14, and A29700G in the 3’UTR region of the virus. The Phe1757Leu modification is located in the middle of a zinc binding site at the interface between the two domains of the Nsp3 protein. It can therefore be surmised that this mutation could subtly change the proofreading process correcting replication errors in a way that would be less amenable to the entry of UTP opposite an A in the negative viral template. We noted that 3 out of the 5 samples that acquired the most C did so through a transition from U to C. The first one, HongKong/HKPU1_2101, shows two simultaneous transitions at positions 12877 and 23857. These mutations being synonymous, they are unlikely to change the replication-correction mechanism. The second one, USA_SC_3571, and the third one, Australia/VIC209, show transitions of the same type, also synonymous, at positions 11635 and 19713 respectively. Finally, the last two samples, USA/WI-33 and USA/WI-28, were derived from the transversion from A to C at position 29567, a mutation at the end of ORF9b.

For dataset number 2, this reversal of the trend concerns mostly Latino-American strains. The succession of mutations C241U, C14408U, then A23403G discussed in relation to the generation of end of translation codons in the virus genes, is followed by C3037U (synonymous), and the triplet G28881A, G28882A, G28883C, overlapping the codons at position 203-204 of the N nucleocapsid N gene. They mutate an arginine-glycine dipeptide into a lysine-arginine dipeptide. This alters the positive charge of the protein and may help improve its role in the assembly of the virus genome in the capsid, as discussed below in relation to the appearance of blooms (36). After this triple modification, we see several reversals of the tendency to lose C in the genome. U29148C (Ile292Thr) is found again in the nucleocapsid N gene, then U27299C (Ile33Thr) in the Orf6 gene, resulting in a set of samples that have at worst gained as much C as they have lost. There are also 3 samples among the 39 in the subtree that gained one more C than they lost (Brazil/RJ-763, Argentina/Heritas_HG007, Argentina/Heritas_HG006). Each time, the last C acquisition comes from a transversion from an adenine (in positions 14164 (Met233Leu), 16076 (Asp870Ala), and 29782, in the late 3’UTR of the viral genome. Overall, it is the change in the nucleocapsid that appears to be most conducive to reversing the tendency to lose C. Indeed, this protein, expressed at a high level during the infection, regulates the process of replication / transcription of the virus and this may account for this remarkable observation [39].

#### Emergence of blooms

The succession C3037U, (C241U, A23403G), C14408U is present upstream of 10 sub-trees, which we considered to be significant (see **Figure 5** and **Materials and Methods**).

**Figure 5:**
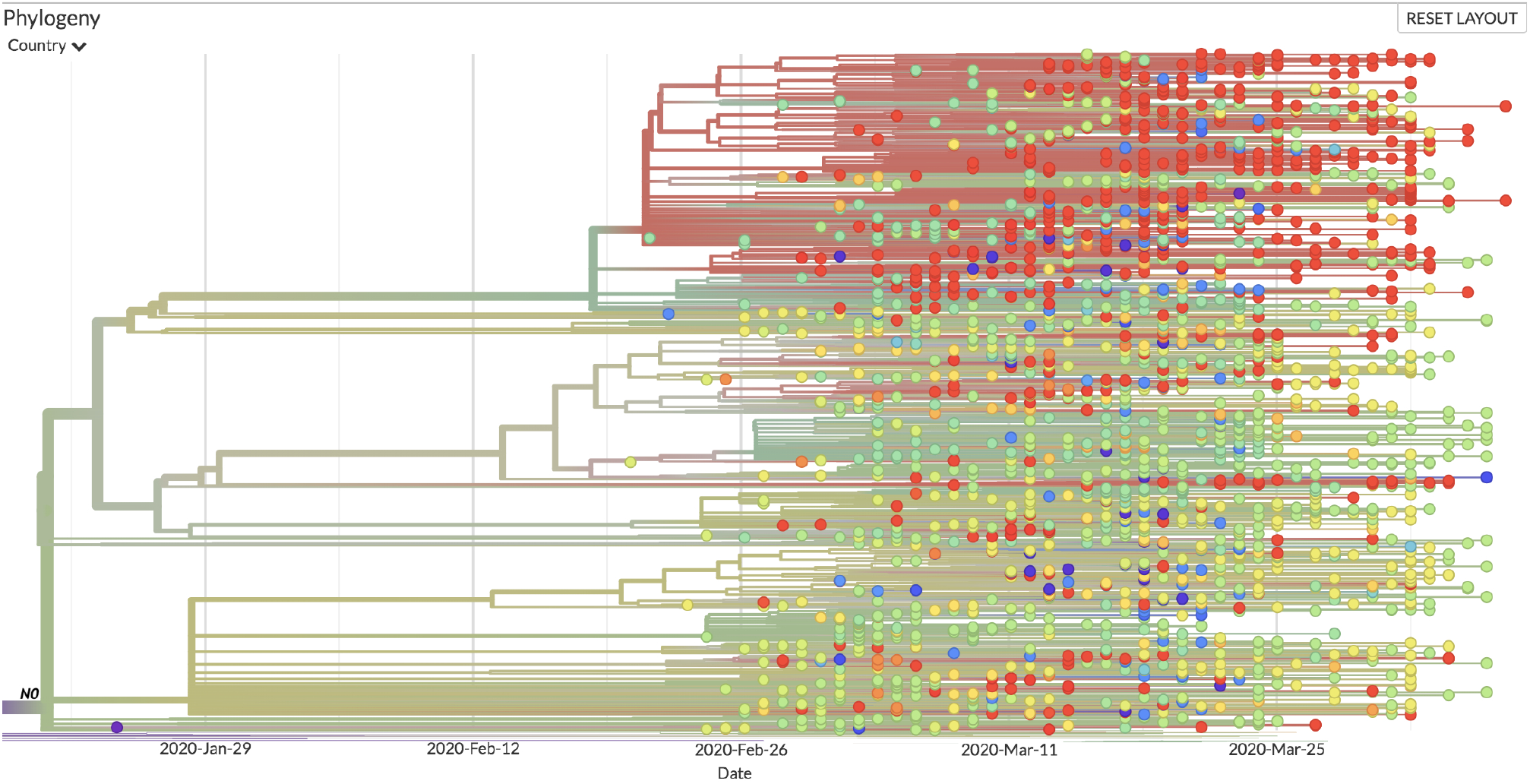
Example of blooms. The subtree shown here contains 10 of the 20 most significant blooms in the sense of the method we used. Node N0 is the place where mutation C14408U emerges.

The synonymous mutation C3037U is located at the end of the ubiquitin-like domain 1 of protein Nsp3. This leads to a sequence (UUUUUU) that promotes changes in the reading frame and could decrease the translation efficiency of the proteins of the ORF1a region. The C241U mutation is observed very often [40]. It is found in the region that initiates replication of the virus. We can therefore assume that this may alter the frequency of replication. A23403G is a widely spread non-synonymous mutation which leads to the replacement of an aspartate by a glycine at position 614 of the spike protein, which is used by the virus to bind its host cell’s receptor. For this reason, several previous analyses have suggested that this mutation has an important role in the spread of the virus [41,42]. Here, the fact that it is part of a major bloom can be considered as an additional argument favouring this interpretation. The C14408U changes an amino acid from proline to leucine (Pro314Leu) just after the end of the NiRAN domain (nidovirus RdRp-associated nucleotidyl transferase) of the protein Nsp12 ending in a “zinc finger”. The NiRAN domain, essential for the replication of the virus, acts as a nucleotidyltransferase, preferring UTP as a substrate for a function which has not yet been clarified [43]. The proline modified in the mutant is part of a dipeptide diproline which plays the role of hinge of separation between the NiRAN domain and the following domain.

A second bloom, which shares several elements with the preceding one begins with the same sequence C3037U, (C241U, A23403G) and C14408U. However it continues with a series of contiguous mutations resulting in a change (G28881A, G28882A, G28883C) in nucleocapsid N, as we saw previously. It is worth noticing that this change might have a role in assembling the virus genome in the capsid by phase separation [38]. This might increase the efficiency of virus transmission and thus contribute to the formation of blooms. The fact that it is a cluster of mutations involving G is intriguing. It may result from the fact that it spans a GGGG sequence.

We have previously seen that mutation G11083U (protein Nsp6, Leu37Phe) has initiated another succession of mutations that led to premature translation termination of a viral protein. Here, this widely distributed mutation is at the root of blooms. As discussed, it is possibly favouring coronavirus infection by compromising delivery of viral components to lysosomes for degradation. This would certainly favour blooms. The mutation is followed, in a first bloom-generating succession, by G26144U (Gly251Val) in protein Orf3a, that forms potassium channels important for innate immunity response - but the exact function of the protein still remains open to question [44]. Subsequent mutations are C14805U (synonymous) and U17247C (synonymous). This succession suggests that the first mutation in protein Nsp6 and perhaps the second one are the primary causes of the bloom [27]. The role of the fist mutation is further substantiated by the second bloomgenerating succession where it is followed by a quadruplet: C6312A (Thr2016Lys) in the interdomain region that precedes domain G2M of multi-domain protease Nsp3, then associated with three C → U mutations, hence expected to be more frequent: C13730U (Ala88Val) in the NiRAN domain of protein Nsp12, C23929U (synonymous), and finally C28311U (in a sequence of four C, Pro13Leu) at the beginning of the nucleocapsid protein, N.

A second succession of mutations that ends up in blooms is C8782U (synonymous), U28144C (Leu84Ser) in protein Orf8, the function of which has been discussed previously and defines a significant clade of the virus variants [24], ending up with C26088U (synonymous). The Leu84Ser mutation co-evolves significantly with the Asp614Gly mutations of the spike protein discussed above [37], which makes it another likely candidate for positive selection leading to increased spreading of the virus, hence blooms.

#### Change in frequency of transitions / transversions

Among the mutations upstream of branches showing significant changes in the transition / transversion flow is the mutation C17747U, which modifies a proline residue into a leucine residue in the protein Nsp13 (**Figure 6** and **Materials and Methods**).

**Figure 6.**
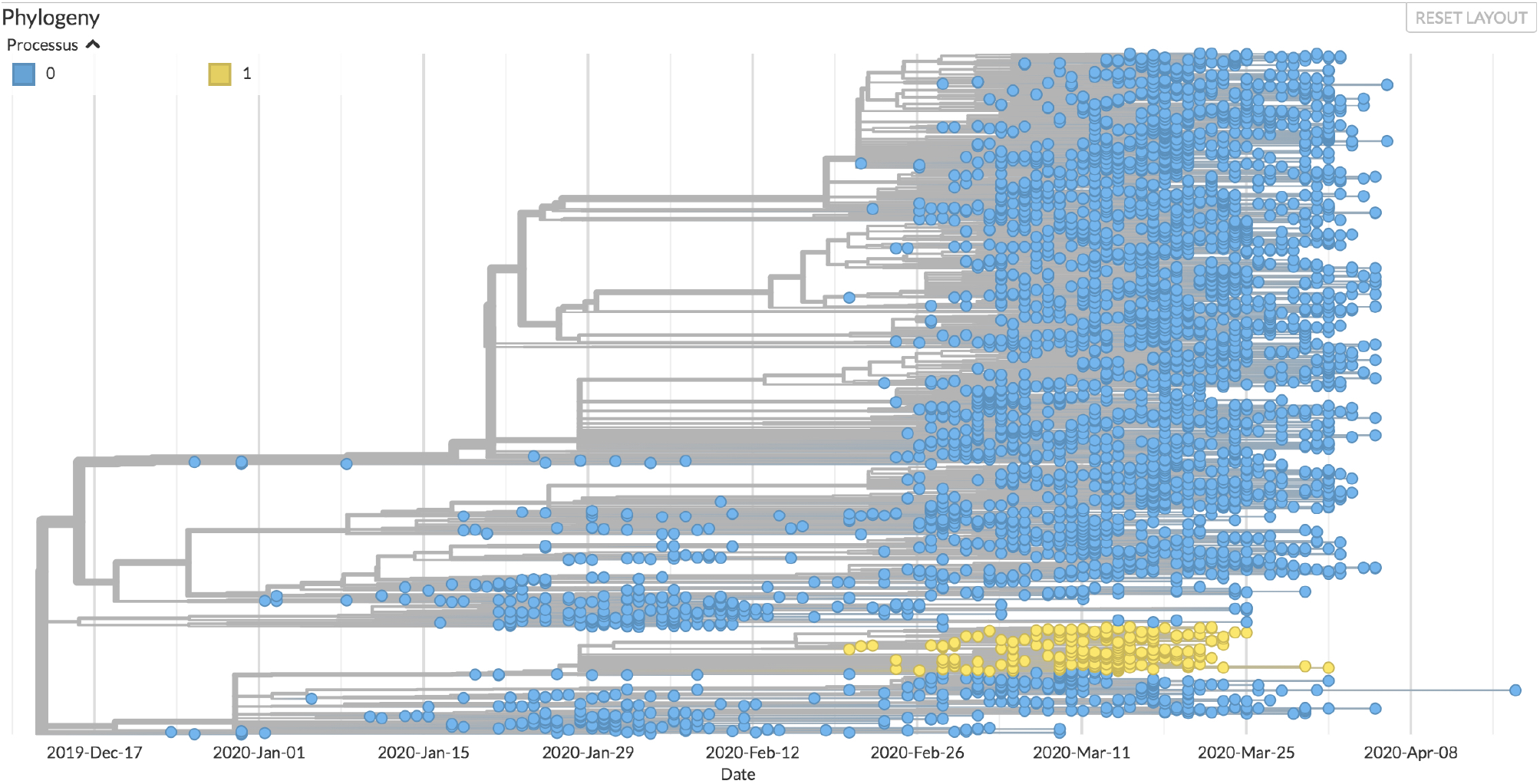
One of the descents considered to be significant for the change in the process of molecular evolution. The progeny resulting from the C17747U mutation is shown in yellow, and its evolutionary process is modelled by a 6-parameter TN93 model (process 1). The rest of the tree (blue leaves) is modelled by a 3-parameter TN93 model.

This mutation affects the protein domain which has nucleoside triphosphatase activity, the exact role of which is unknown but consistent with a proofreading activity [45]. We might propose that it is involved in the quality control of the product of the replication of the virus for example *via* stabilizing the “anti” form of nucleotides, thus avoiding the mismatching leading to transversions. In fact, this protein has been identified among those which lead to a significant alteration in the diversity of the viral genome [16]. The existence of a notable change in the type of mutations located downstream of the tree is therefore a strong argument for the discriminating role of the corresponding region of the protein. Furthermore, to the extent that this mutation increases the frequency of mutations in a biased manner, we can expect the ensuing descent to lead to an attenuation of the virus. However, as this changes the evolutionary landscape, this evolution could lead to “innovative” mutations modifying the pathogenicity of the virus, and this especially under conditions where recombination due to co-infections would be favoured. This is yet another argument for choosing a strong public health policy which tends to avoid the formation of clusters of infection.

### Conclusions and perspectives

The COVID-19 epidemic is a life-size experiment in virus evolution. Remarkably, we neither know the real origin of the virus [46], nor where it will lead us. This explains why the vast majority of studies of the SARS-CoV-2 virus and its evolution are essentially descriptive. Here, we tried to make use of the ongoing evolution of the virus to investigate some of its related constraints using a hypothesis-driven probabilistic modelling approach to the molecular evolution of the virus. Based on the assumption that the virus’metabolism is ruled by its host. Based on the metabolic set up of the host cells, acting as a compulsory material framework for the multiplication of viral particles, we pointed out specific changes in the evolution pattern of the virus descent, witnessed by changes in the virus genome composition as time passes. Using the widely spread C to U change in this genome’s composition as a base line, we identified nodes where the change is shifted from this direction to another one, favouring transversions rather than transitions, reversing the C to U trend towards U to C enrichment or generating blooms with sudden appearance of multiple branches in the evolution tree. This allowed us to point out a series of functions that are evolving towards a more efficient spread of the virus (e.g. the previously identified Asp214Gly mutation of the spike protein, but also the Gln57His mutation of the Orf3a potassium channel). We also noticed that Orf8 is the likely site of an ongoing competition for expression of two frameshift-dependent overlapping proteins Orf8a and Orf8b. Similarly, the unstable region of Orf7 could promote the synthesis of the very small membrane protein Orf7b, whose function remains unknown to date. Finally, the reversion of the tendency to favour U over C indicates that nucleocapsid protein N may be involved in the control of CTP synthesis in the host, suggesting an interesting target for future control of the virus development. We hope that this combination of mathematical and biochemical knowledge will help us devise further enterprises against the dire consequences of COVID-19. We noticed that among the possible way for the virus to escape CTP-dependent control in cells would be to infect cells that are not expected to grow, such as neurons. This may account for unexpected body sites of viral development observed in the present epidemic.

## Acknowledgements

AL would like to thank the Centre Interdisciplinaire de Recherche en Biologie (CIRB, Collège de France) for its funding, as well as the members of the SMILE (Stochastic Models for the Inference of Life Evolution) team of the CIRB for many fruitful discussions on the modelling of the COVID-19 epidemic. AD thanks Stellate Therapeutics for the support of his laboratory.

## Materials and Methods

### Data processing

A total of 4,792 sequences of the SARS-CoV-2 virus were recovered from the GISAID databank [47] on April 21, 2020 for the first dataset. Only the genomes of viruses from the human hosts of SARS-CoV-2 of a length greater than 25,000 bp were retained. Sequences for which the sampling date was insufficiently informed (absence of the harvest day, sometimes of the month) were also excluded. For sequences present multiple times, only the first isolate was retained. We also reused the work of the Nextstrain teams and discarded the too divergent or unstable samples that they themselves had left out (github.com/nextstrain/ncov/blob/master/defaults/exclude.txt).The sequence of 26 coding regions (Nsp1, Nsp2, Nsp3, Nsp4, Nsp5, Nsp6, Nsp7, Nsp8, Nsp9, Nsp10, Nsp11, Nsp12, Nsp13, Nsp14, Nsp15, Nsp16, S, ORF3a, E, M, ORF6, ORF7a, ORF7b, ORF8, N and ORF10) was characterized using NC_045512 as a reference. The total number of sequences retained at the end of the treatment is 4,088 sequences. A second dataset of 3,246 sequences, 510 of which are common with the first dataset, was retrieved on July 6, 2020 using directly the Nextstrain API [48].

We note here that, over time, data availability kept being altered, with some sequences deleted from the samples, while other ones entered the database. Furthermore, it was generally difficult to extract large samples of sequences so that it was extremely difficult to build up a consistent data repository where correct statistical approaches could be implemented. It seems very awkward that the bulk of the sequences of a virus of worldwide importance has not been made available at the International Nucleotide Sequence Database despite recommendations of the major research institutions [49,50].

### Phylogenetic reconstruction

The reconstruction process begins with aligning all of the sequences to the reference sequence. Insertions and deletions of genome regions were not taken into account. It was out of the question here to take into account nucleotide insertions and deletions. We retained only the potential one to one substitutions. We used program MAFFT [51] to generate these alignments. Some ambiguous positions were highlighted during the alignment process. For example, some regions of the genome may display high instability and wide variability depending on the parameters of the algorithm used to perform the alignment. To overcome this problem, we used the same masks as those used by the Nextstrain team. Sites 18529, 29849, 29851, 29853, as well as the first 130 and last 50 sites of the genome were therefore omitted from the substitution analysis. We used a General Time Reversible (GTR) model to infer the substitution process at work using the IQTREE software [52]. This first tree is a fairly raw version which does not take into account the temporal aspect of evolution. The Treetime software [53] allows you to refine this tree by also taking into account the sampling dates of the sequences. It then reconstructs the tree with maximum likelihood compared to the sampled sequences. Using maximum likelihood approaches it also infers the compositions of the ancestral sequences of the samples, as well as a 90% confidence interval around the most likely date of these common ancestors. Once the tree has been created, we could then reconstruct the order in which the mutations in each sample appeared, in the sense of maximum likelihood. For the visualization of the tree and the production of Figures 2 to 6, we used the Auspice program developed by Nextstrain, to which we made some modifications to display the parameters we were interested in. To this purpose, we developed a Python script to modify the JSON file used as input by the Auspice program. This allowed us to enrich the visualization capabilities of the software by adding quantities such as the number of C acquired or lost by a sample compared to the reference and to generate original tree presentations.

### Identification of blooms

The main pitfall we had to face when identifying blooms was the bias introduced when selecting samples from the phylogenetic tree. In particular, some hospitals were likely to provide more samples than others, due to the different health policies and means implemented depending on the country. In order to avoid selecting nodes likely to generate a bloom due to oversampling, we chose to develop a custom-made statistical method meant to cope with this difficulty.

A subtree is any set of nodes and leaves rooted in one of the nodes of the main tree. The idea is to use the information provided by the identity of the countries represented in each sub-tree: the easier a strain is spread, the higher the number of countries in which it is expected to be observed. To implement this heuristic, it is necessary to control two factors: the size of the tree (two trees of unequal depth, that is to say rooted on different dates, naturally show diversity as different countries) and the heterogeneity of sampling (countries where sampling and sequencing are carried out with different intensities have different probabilities of appearing in a given sub-tree).

These two factors interact, because the size of a tree (the number of its leaves for example) obviously varies with the sampling intensity. One way to control this interaction is to measure the size of a tree by its total length, or sum of branch lengths, in time units. Indeed, this observable is not very sensitive to the effects of oversampling because the presence of many sequences sampled in the same place at about the same time generates a sub-tree whose length is close to zero.

To control the effect of the length factor L on the number of countries represented, N, we seeked to learn the relation N = f(L) in a typical tree in order to be subsequently able to identify the sub-trees whose number of countries represented, for a known length L, exceeds the expected f(L). A simple statistical model consists in supposing that the number of occurrences of country i in a tree of length L is a Poisson distribution of parameter θ_i_L and that these numbers are independent. If K is the total number of countries referenced by Nextstrain, the number of countries N represented in a tree of length L is therefore the sum of K Bernoulli variables independent of parameters 1 - exp (- θ_i_L). For example, if countries are divided into two groups, the k_1_ ‘frequent’ of intensity θ_1_, and the k_2_ ‘rare’ of intensity θ_2_≪θ_1_, N has the mean K – k_1_exp(-θ_1_L) – k_2_exp(-θ_2_L),

which behaves when L is large like K – k_2_exp(-θ_2_L).

In addition, when L is large, assuming that θ_2_L = O(1), the distribution of N is approximately equal to k_1_ + N_2_, where N_2_ foloows a Poisson law of parameter k_2_(1-exp(-θ_2_L)).

So, we used the parameterization:

N = a - bexp(-cL), interpreting the parameters as follows: a is the maximum number of countries, b is the number of countries with low sampling/sequencing intensities and c is a density of presence of these countries per unit length of tree. Under the null hypothesis, N is distributed as a - b + N_1_, where N_1_, follows the Poisson’s law of parameter b(1-exp(-c L)). Finally, we selected the 20 most significant blooms, i.e., those whose behaviour deviated the most from that expected by our estimator. This then allowed us to reconstruct the lineages and the mutations that appeared successively upstream of each node at which a bloom had occurred. This allowed us to identify the succession of mutations common to some of these nodes and thus those giving rise to the majority of statistically significant blooms. Furthermore, we restricted the automatic selection of nodes so that no selected node was present in the lineage of another one. The selected blooms are therefore mutually independent, even though they may obviously have common ancestors. To arbitrate the choice between two nodes present in the same lineage, we have systematically kept the oldest node, and thus the most dense tree.

### Detection of changes in the molecular evolutionary process

We investigated whether the substitution process in some sub-trees behaved differently from what was observed in the rest of the tree, statistically speaking. To this aim, we used the classical TN93 model from Tamura and Nei [54] with 3 parameters (purine transition rate, pyrimidine transition rate and transversion rate) and allowed these three rates to take, downstream of a candidate node Ni, values that differed from those they take in the rest of the tree. We then used a second (6-parameter) model. Since this model is nested in the first (3-parameter) model, we used as test statistic the likelihood ratio 2Δl = 2(l_1_ - l_0_), where l_0_ is the log-likelihood under assumption H0 (3-parameter TN93 model estimating all the elements of the tree) and 1_1_ is the log-likelihood under assumption H_1_ (6-parameter TN93 model with local differentiation of the parameters downstream of a node of interest). We then compared the likelihood ratio to a distribution of the \chi^2^ with 3 degrees of freedom, whose significance threshold at 5% is 7.81. We were then able to identify the nodes from which the evolution process varied significantly and to quantify the variations of the different substitution rates, i.e. the nodes for which we can reject the H0 hypothesis that the 3-parameter model TN93 produces better estimates of tree substitution rates than the 6-parameter model TN93. We have chosen to implement these models ourselves in Python, in order to keep this parametrization flexibility. The program allows us to determine the set of nodes and leaves present downstream of a node of interest and to perform hypothesis testing by calculating the likelihood ratio and the different substitution rates.

#### TEXT BOX

##### Molecular evolution and Muller’s Ratchet

Biology rests on the laws of physics. Because it develops at approximately 300K, it is subject to the universal stress of thermal noise, involving an energy that does not differ considerably from that involved in the chemical bonds of biological chemistry. It follows that the reactions that come through and organize living things cannot develop with strict reproducibility. Inevitable errors cause the product of a reaction to differ from what it is supposed to be. Genome replication cannot escape this constraint. The consequence is that, in the progeny of a virus, there is always a number of variants, named mutants when they carry over alterations of the genome. In most cases, these mutants correspond to the change from one of the four nucleotides to a different one. This process, as a rough approximation, is random — the mutant position can be anywhere in the genome, and the replacement of one nucleotide is by any of the other three. As time goes by, all the nucleotides of the genome are likely to change into others. This will affect the functions necessary for the multiplication of the virus, and some changes will continue to be propagated (be fixated), while others will end up without a progeny. A mutation followed by a fixation is called a substitution. The substitution of a purine for a pyrimidine (or vice versa) is called a transversion; other substitutions are called transitions. The likelihood of a particular mutation returning to the ancestral state is very low. The probability that a particular mutation will return to the ancestral state is very low. This forces evolution to always go forward, without the possibility of going back. This process was noticed in 1932 by Hermann Muller in the special case of the effects of irradiation on mutagenesis. His reflection has since been simplified and popularized. It is now known as the “Muller’s ratchet” [55]. It is obviously highly probable that the majority of mutations leads to the partial or total loss of the functions coded by the altered regions of the genome. It follows that this generally leads, in the long term - but not in the short term - to the attenuation of the functions allowing multiplication and virulence of pathogenic species. This is why Louis Pasteur and his successors could have the luck to isolate attenuated organisms which, in some - rare - cases, could then be used for the vaccination of infected persons [56]. However, this process becomes unproductive as soon as co-infection with different mutants occurs under circumstances where recombination is possible. Two different mutants can recombine into the ancestral form of the pathogen and erase the entire benefit of attenuation. This is all the more harmful since the old forms are also, very often, those which spread most easily.

**Boxed text Figure.**
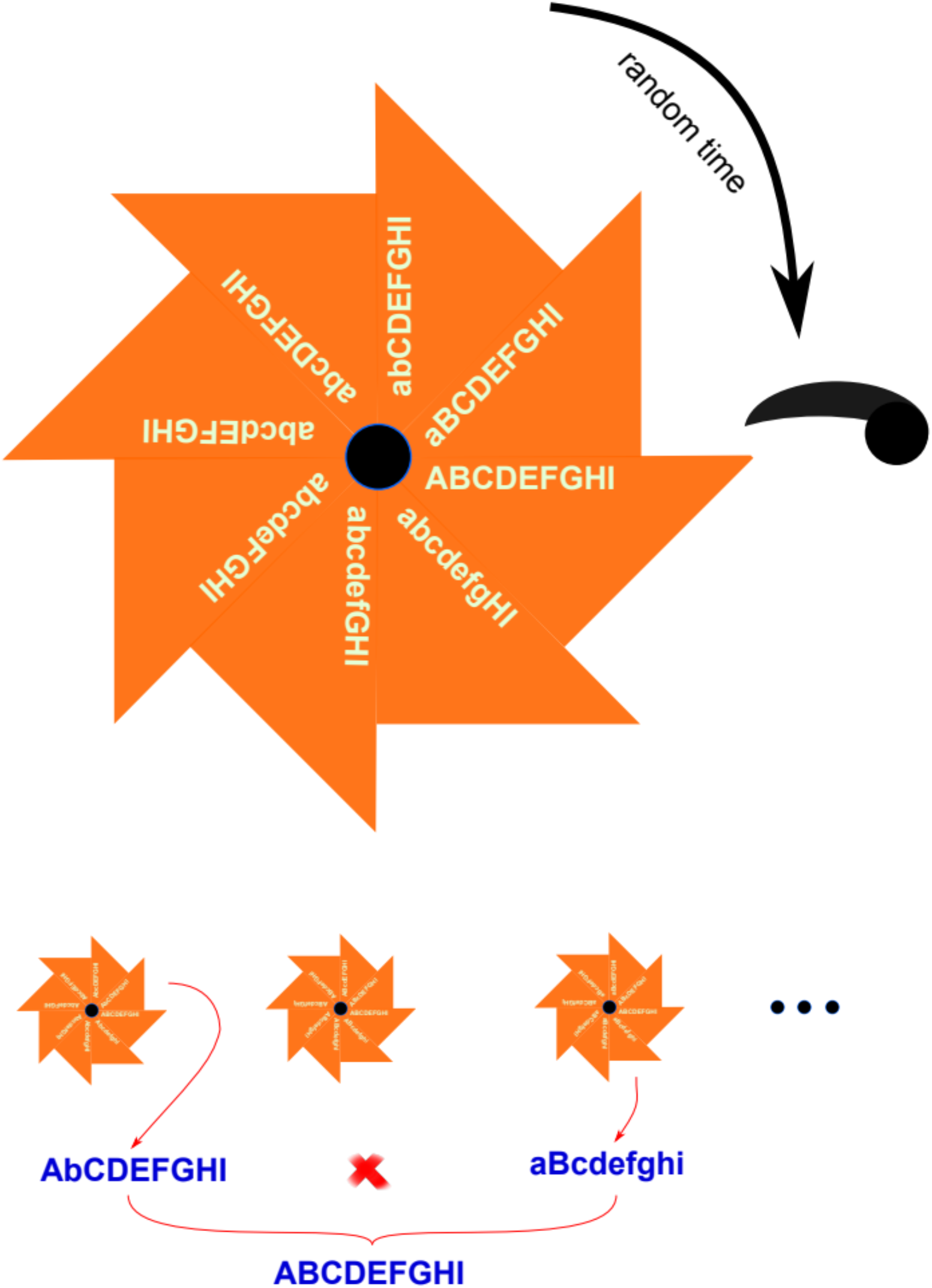
Muller’s ratchet and recombination. The figure is reprinted from reference [57]. Genes (capitals) are mutated at random in a different form (low case). Mutations accumulate ratchet-like because the probability of reversion to the parent form is negligible. This happens independently for viruses of different descents. However, if viruses from different descent happen to be in the same cell, they can recombine. This allows them to recreate the ancestral form of the virus.

